# Astrocytes dystrophy in ageing brain parallels impaired synaptic plasticity

**DOI:** 10.1101/2020.08.05.237420

**Authors:** Alexander Popov, Alexey Brazhe, Pavel Denisov, Oksana Sutyagina, Natalia Lazareva, Alexei Verkhratsky, Alexey Semyanov

## Abstract

Little is known about age-dependent changes in structure and function of astrocytes and of the impact of these into the cognitive decline in the senescent brain. The prevalent view on age-dependent increase in reactive astrogliosis and astrocytic hypertrophy requires scrutiny and detailed analysis. Using two-photon microscopy in conjunction with 3D reconstruction, Sholl and volume fraction analysis we demonstrate a significant reduction in the number and the length of astrocytic processes, in astrocytic territorial domains and in astrocyte-to-astrocyte coupling in the aged brain. Probing physiology of astrocytes with patch-clamp and Ca^2+^ imaging revealed deficits in K^+^ and glutamate clearance, and spatiotemporal reorganization of Ca^2+^ events in old astrocytes. These changes paralleled impaired synaptic long-term potentiation (LTP) in hippocampal CA1 in old mice. Our findings may explain astroglial mechanisms of age-dependent decline in learning and memory.

## 1. Introduction

The ageing of the brain represents the life-long adaptation of this exceptionally complex organ to the environmental “exposome” (cumulative exposure of an individual to environmental factors during the lifetime) interacting with the genetic background. This adaptation, in the form of much-praised neuroplasticity, proposed over a century ago by Willaim James (James, 1890) and conceptualised by Eugenio Tanzi and Ernesto Lugaro (Lugaro, 1898; Tanzi, 1893), continuously remodels neuronal networks thus tuning the brain structure and function to match continually changing milieu. This neuroplasticity is achieved by the close coordination in neuronal and glial networks, in which glia act as a principal homeostatic element supporting neuronal function, preserving the integrity of neuronal ensembles and connectome, assisting in synaptogenesis and defending the brain against insults. Astrocytes, in particular, control homeostasis of ions and neurotransmitters, regulate brain blood pressure, thus containing excitotoxicity and defend the brain against oxidative stress associated with high energy consumption of neurons (Marina et al., 2020; Verkhratsky and Nedergaard, 2018).

Astrocytic ageing remains rather poorly characterised; neither their structure nor function have been analysed in depth. It is generally agreed, based on stereological and immunohistochemical studies, that the total numbers of astrocytes in the central nervous system (CNS) of rodents, primates and humans does not significantly change with age (Fabricius et al., 2013; Olabarria et al., 2010; Pelvig et al., 2008; Robillard et al., 2016). Early investigations frequently described an increase in CNS expression of glial fibrillary acidic protein GFAP in rodents (Goss et al., 1991; Kohama et al., 1995; Nichols et al., 1993) and humans (David et al., 1997). Morphological analysis of aged astrocytes revealed controversial results. Increase (Hayakawa et al., 2007), decrease (Cerbai et al., 2012) or no change (Robillard et al., 2016) in the number of GFAP-positive astrocytes has been reported. Similarly, both hypertrophy and atrophy of GFAP-positive were observed, often with certain region specificity (Bjorklund et al., 1985; Cerbai et al., 2012; Rodriguez et al., 2014). Labelling of astrocytes with other markers such as antibodies against glutamine synthetase or S100B demonstrated minor (and again region-specific) age-dependent changes in cellular morphology (Rodriguez et al., 2014). The lifelong analysis demonstrates that astrocytes undergo biphasic morphological changes – the astrocytic domain and complexity increases from juvenile to middle age, with subsequent decreases in the old age (Robillard et al., 2016).

There is not much known about changes in astrocytic physiology in ageing. The main biophysical properties of the plasmalemma seem to change very little; the resting membrane potential and input membrane resistance do not differ between young and old rodents (Lalo et al., 2011). Astrocytes from old animals generate membrane currents and Ca^2+^ responses to major neurotransmitters, indicative of expression of ionotropic glutamate and P2X receptors as well as metabotropic glutamate, noradrenaline, cannabinoid and P2Y receptors (Gomez-Gonzalo et al., 2017; Lalo et al., 2018; Lalo et al., 2011). Similarly, astrocytes seem to maintain functional glutamate transporters (Lalo et al., 2011) responsible for glutamate clearance.

Brain ageing and neurodegeneration are universally regarded as synaptic dysfunction. Astrocytes perisynaptic processes cover ~50 – 60% of all synapses in the CNS, with particularly high coverage of complex excitatory synapses (Gavrilov et al., 2018; Patrushev et al., 2013). The thin leaflets of astrocytic membranes that enwrap synaptic structures form the astrocytic synaptic cradle, which controls synaptogenesis, synaptic maintenance and synaptic elimination (Verkhratsky and Nedergaard, 2014). Astrocytic perisynaptic processes are dynamic structures, which by changing their morphology affect the degree of synaptic coverage thus influencing synaptic transmission. Such morphological plasticity of astrocytes, in particular, occurs upon systemic states (such as lactation (Oliet et al., 2001)) or in response to environmental stimulation (such as dietary changes (Popov et al., 2020)). Classical analysis of astrocytic morphology based on GFAP immunocytochemistry does not access perisynaptic structures because of their small size and absence of GFAP filaments. Here we studied the morphology of astrocytes in hippocampal slices of young, adult and old mice using two-photon imaging and volume fraction analysis. We correlated age-dependent morphological changes with astrocytic physiology. Our results demonstrate a decrease in astrocytic domain size, volume fraction of perisynaptic processes, and astrocyte coupling through gap-junctions in the ageing; this, in turn, affects K^+^ buffering and glutamate clearance by astrocytes, and astrocytic Ca^2+^ signalling, which all impact on synaptic plasticity and affect long-term potentiation in hippocampal synapses.

## 2. Results

### 2.1. Biphasic morphological changes of astrocytes during the lifespan

Mice of three age groups were used in this study: young (3-4 months old), adult (9-12 months old) and old (20-24 months old). These ages roughly correspond to human ages of 20 - 30 years, 38-47 years and 56 - 69 years, respectively (Flurkey et al., 2007). Astrocytes in CA1 *stratum (str.) radiatum* of hippocampal slices were loaded with fluorescent dye Alexa Fluor 594 through a patch pipette to visualise their morphological profiles and analyse their processes. Astrocytic processes are morphologically and functionally classified into three classes: (i) branches and branchlets - main processes containing organelles, (ii) leaflets - flat organelle free peripheral and perisynaptic processes and (iii) endfeet covering the blood vessels (Patrushev et al., 2013; Semyanov, 2019). The Z-stacks of images, taken with two-photon imaging, were used for three-dimensional (3D) reconstruction of astrocyte soma and branches (Fig. 1a) and subsequent Sholl analysis. We quantified the number of intersections of branches with the concentric spheres of increasing radius plotted around the centre of astrocyte soma (Supplementary Fig. 1 and Fig. 1b).

**Figure 1.**
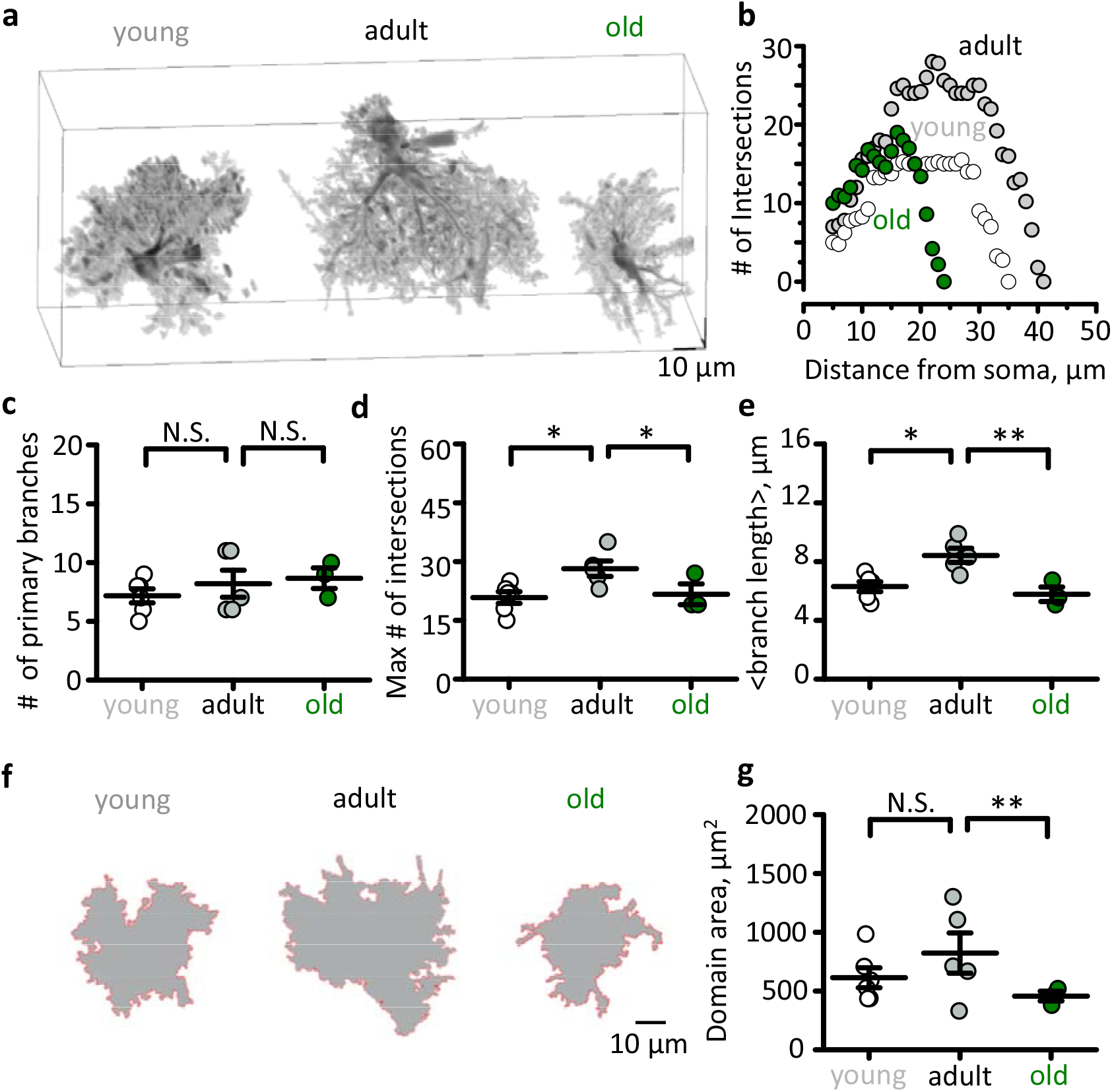
Bidirectional changes in astrocytic branches and branchlets, and of astrocytic domain size during mouse lifespan. **a.** Three-dimensional reconstructions of hippocampal astrocytes loaded with a fluorescent dye (Alexa Fluor 594) of three age groups young (*left*), adult (*middle*) and old (*right*). **b.** An example of single astrocyte 3D Sholl-analysis which shows the number of intersections of astrocytic branches and branchlets with concentric spheres centred in the middle of cell soma. **c-e.** Summary of the primary brunches number (**c**), the maximum number of intersections (**d**) and mean branch and branchlet length (**e**) in three age groups. **f**. Projections of astrocytic reconstructions which were used to define astrocytic domain area in three age groups. **g.** Summary of the astrocytic domain areas in three age groups. *White circles –* young*, grey circles –* adult *and green circles –* old mice. The data are presented as the mean ± SEM. N.S. *p* > 0.05; \**p* < 0.05; \*\**p* < 0.01; two-tailed two-sample *t*-test.

The number of primary astrocytic branches was not significantly different between astrocytes from young, adult and old animals (7.2 ± 0.6, *n* = 6, in the young; 8.2 ± 1.2, *n* = 5, in the adult; 8.7 ± 0.9, *n* = 3, in the old; *p* = 0.21 between young and adult, *p* = 0.79, between adult and old, two-sample *t*-test; Fig. 1c). However, both the maximal number of intersections (20.8 ± 1.9, *n* = 6, in the young; 28.2 ± 1.9, *n* = 5, in the adult; 21.6 ± 2.6, *n* = 3, in the old; *p* = 0.014 between young and adult, *p* = 0.04 between adult and old, two-sample *t*-test; Fig. 1d) and the mean branch length (6.3 ± 0.3 μm, *n* = 6, in the young; 8.4 ± 0.5 μm, *n* = 5, in the adult; 5.7 ± 0.5 μm, *n* = 3, in the old; *p* = 0.005 between young and adult, *p* = 0.005 between adult and old, two-sample *t*-test; Fig. 1e), were significantly larger in the adult than in the young or in the old mice. Such biphasic age-dependent changes of astrocytic branches impacted upon the size of an astrocytic domain which was measured as the area of the maximal projection of an astrocyte along Z-axis (Fig. 1f). Indeed, the domain area was the largest in astrocytes of the adult as compared to either young or old mice. However, the difference reached statistical significance only between adult and old mice (613 ± 85 μm^2^, *n* = 6, in the young; 823 ± 171 μm^2^, *n* = 5, in the adult; 457 ± 41 μm^2^, *n* = 3, in the old; *p* = 0.28 between young and adult, *p* = 0.049 between adult and old, two-sample *t*-test; Fig. 1g). These findings are consistent with a previous report that astrocytic structure becomes more complex and size increases during development to the adulthood, then the astrocytes shrink with ageing (Robillard et al., 2016). We are interested in the morphofunctional changes that occur in astrocytes in the old age; hence, thereafter, we focused on the comparison of adult and old mice astrocytes.

### 2.2. Decreased volume fraction (VF) of fine astrocytic processes in the aged brain

The above analysis reports change only in primary astrocytic processes, the branches and branchlets that are thick enough to be identified in two-photon fluorescence images. However, a mass of tiny peripheral astrocytic processes, the leaflets, cannot be resolved with diffraction-limited optical methods (Gavrilov et al., 2018). To quantify age-related changes in peripheral and perisynaptic processes, we analysed their VF as previously described (Medvedev et al., 2014; Popov et al., 2020; Wu et al., 2019). The VF of optically unresolved processes was estimated as a ratio of fluorescence along the astrocytic anatomic domain cross-section to the maximal fluorescence of soma (Fig. 2a,b). To reduce selection bias, five cross-sections were done for each cell. Then the fluorescence peaks corresponding to branches and branchlets were removed. Two VF values were obtained for each cross-section: one on each side of soma. Mean VF was significantly reduced in the old animals (4.5 ± 0.4 %, *n* = 50 (5 slices) in the adult; 2.1 ± 0.2 %, *n* = 30 (3 slices) in the old; *p* < 0.001, two-sample *t*-test; Fig. 2c) indicating atrophy of fine astrocytic processes. Notably, the astrocyte shrinkage was not accompanied by a reduction in the overall tissue density of astrocytes bulk-stained with astrocyte-specific marker sulforhodamine 101 in hippocampal CA1 *str. radiatum* (0.59 ± 0.05 astrocytes per 100 x 100 μm^2^, *n* = 10, in the adult; 0.61 ± 0.04 astrocytes per 100 x 100 μm^2^, *n* = 7, in the old; *p* = 0.87, two-sample *t*-test; Fig. 2d). This suggests that astrocyte borders move away from each other which may affect intercellular connectivity through gap-junctions. Because gap-junctions are permeable for small molecules such as fluorescent dyes, we counted the number of astrocytes stained with Alexa 594 by its diffusion through gap-junctions from the astrocyte patch-pipette loaded with the dye (Fig. 2e) and found that the number of coupled cells decreased with ageing (6.8 ± 0.9, *n* = 5, in the adult; 3.0 ± 0.5, *n* = 3, in the old; *p* = 0.015, two-sample *t*-test; Fig. 2e,f).

**Figure 2.**
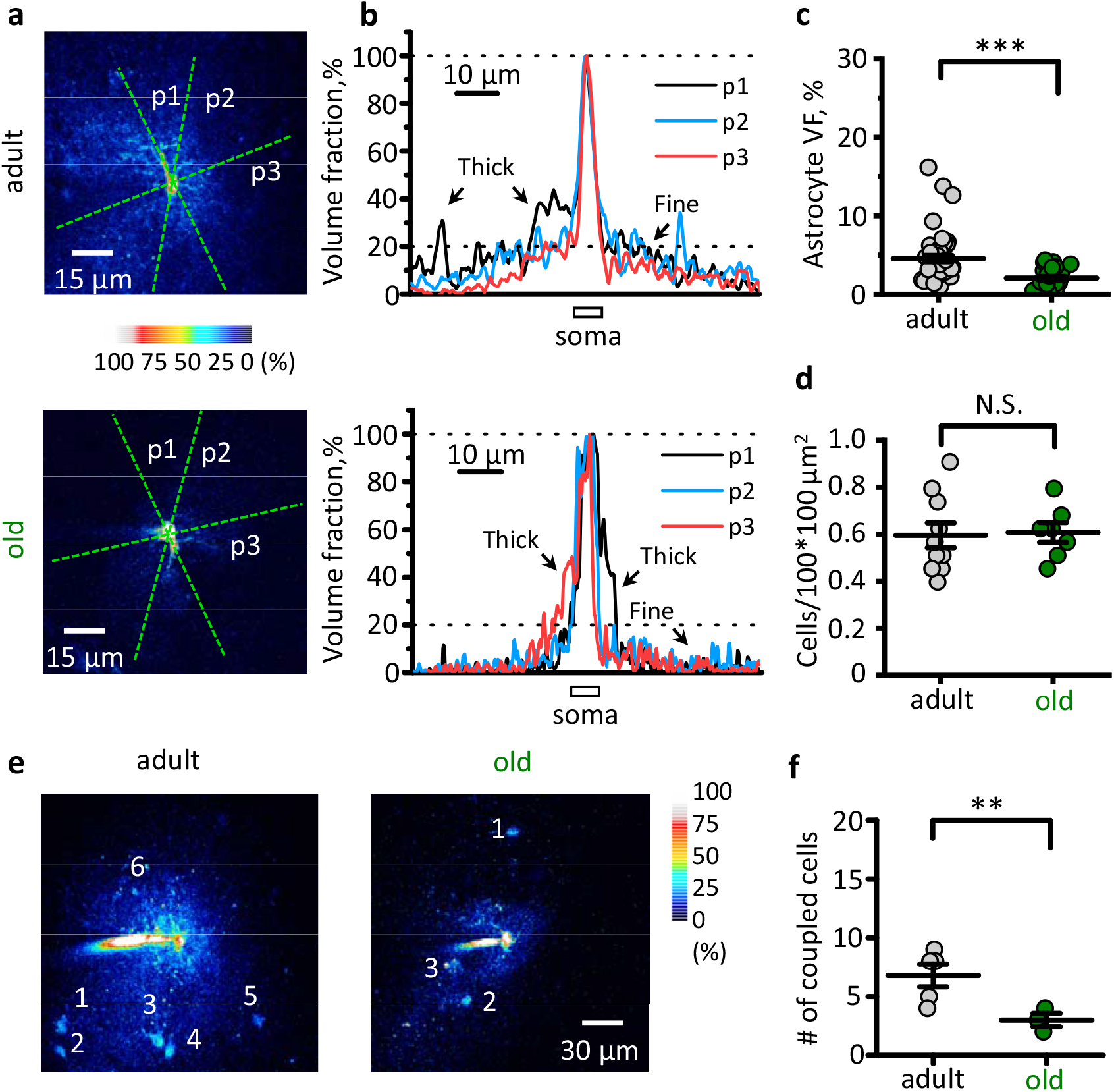
Ageing decreases the volume fraction (VF) of fine astrocytic processes and astrocytic coupling to the neighbours. **a.** Examples of two astrocytes from adult (*top*) and old (*bottom*) mice. The astrocytes were loaded with Alexa Fluor 594 through patch pipette. The fluorescence intensity was of each pixel was normalised to the peak fluorescence intensity in the soma (100 % of VF). Green dashed lines indicate several positions of cross-sections (p1, p2, p3). **b.** Fluorescence profiles corresponding to cross-sections shown at the panel **a**. The central peak of fluorescence corresponds to the brightest pixels in soma to which the fluorescence was normalised to obtain local VF. Other peaks of fluorescence correspond to thick astrocytic branches which were excluded from further analysis. *Top* adult mouse; *bottom* – old mouse. **c.** Summary of fine unresolved processes VF. **d.** The density of astrocytes stained with astrocyte-specific marker sulforhodamine 101. **e.** Examples of dye diffusion through gap-junctions to neighbouring astrocytes from adult (*left*) and old (*right*) mice. *Grey circles –* adult *and green circles –* old mice. The data are presented as the mean ± SEM. N.S. *p* > 0.05; \*\**p* < 0.01; \*\*\**p* < 0.001; two-tailed two-sample *t*-test.

### 2.3. Membrane properties of old astrocytes

Astrocyte morphological remodelling can affect various aspects of astrocyte physiology such as membrane ionic permeability, potassium and glutamate clearance and Ca^2+^ dynamics (Plata et al., 2018; Popov et al., 2020). Although astrocytes are electrically non-excitable cells and do not generate action potentials, they are not electrically idle. Increases in extracellular K^+^ that accompany neuronal activity and synaptic transmission depolarise astrocyte membrane and may affect neurotransmitter transporters (Kirischuk et al., 2016; Lebedeva et al., 2018). Hence, the cell input resistance (R_in_) is an important parameter that determines the astrocyte response to extracellular K^+^. Ageing-dependent astrocyte shrinkage, as well as gap-junction uncoupling, can both increase the cell input resistance (R_i_) (Adermark and Lovinger, 2008). First, we obtained current-voltage relationships (I-V curves) of astrocytes by applying voltage steps of a different magnitude to voltage-clamped astrocytes (Fig. 3a). The I-V curves were linear in astrocytes from both age groups, indicating passive astrocyte properties. However, the I-V curve slope decreased in the older animals pointing to higher cell R_i_ (Fig. 3b) Indeed, R_i_ significantly increased in the old mice astrocytes (21.5 ± 4.5 MOhm, *n* = 8, in the adult; 35.8 ± 2.4 MOhm, *n* = 8, in the old; *p* = 0.007, two-sample *t*-test; Fig. 3c).

**Figure 3.**
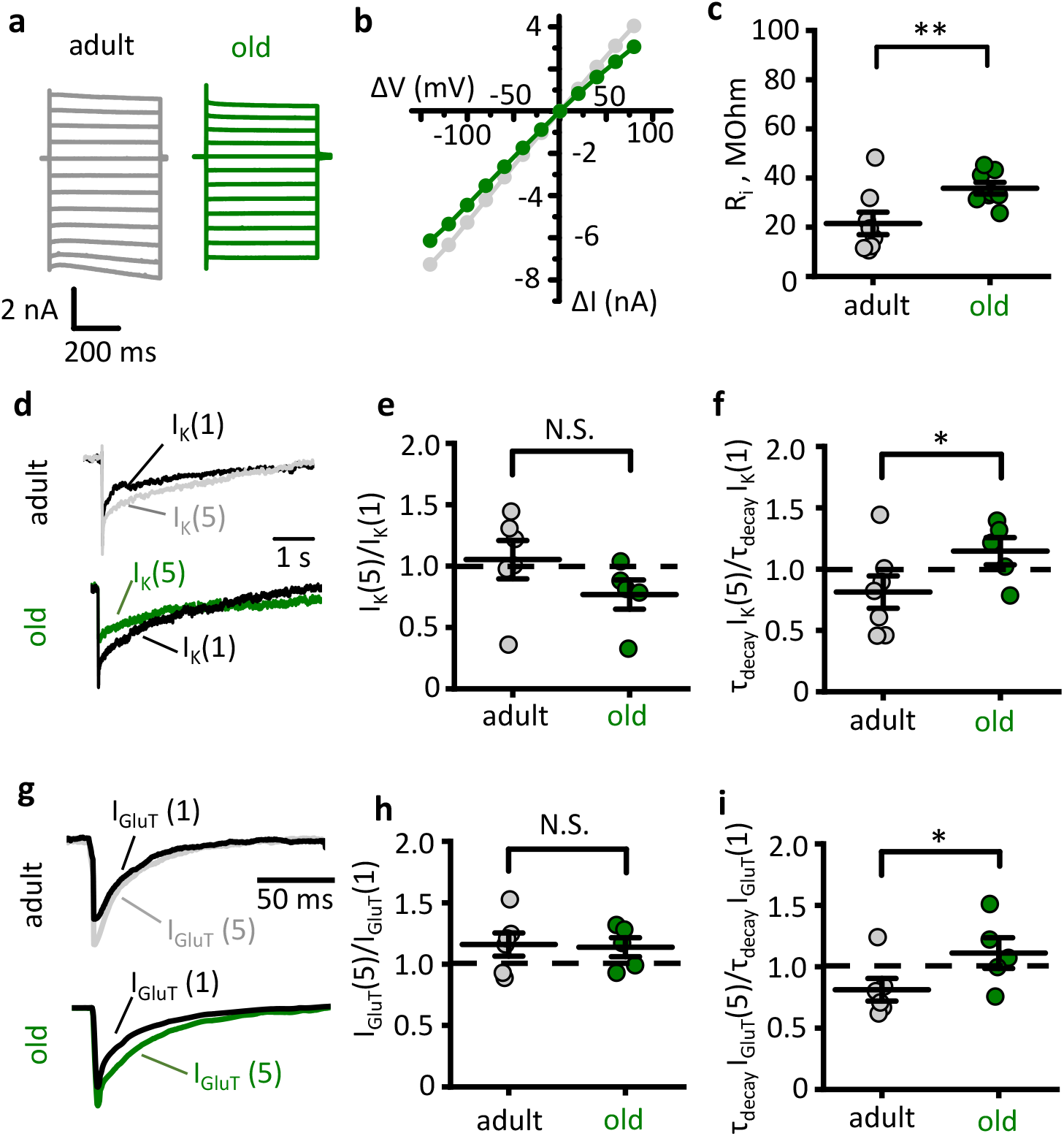
Astrocyte dystrophy enhances glutamate and potassium spillover. **a.** Recordings of astrocytic currents in response to voltage steps (from −140 mV to +80 mV with 20 mV interval) delivered through patch pipette to voltage-clamped adult (grey traces) and old (green traces) astrocytes. **b.** Current-voltage (I-V) relationships based on responses presented on panel a. ΔV – voltage step amplitude, ΔI – current response. **c.** Summary of astrocyte input resistance (R_i_). **d.** Astrocytic currents in response to stimulation of Schaffer collaterals: I_K_(1) – response to a single stimulus; I_K_(5) – isolated response to 5^th^ stimulus in burst stimulation (5 stim x 50 Hz minus 5 stim x 50 Hz). *Top* adult, *bottom* – old mouse. **e,f.** Summary of activity-dependent change in the amplitude [I_K_(1)/I_K_(5), **e**] and the decay time of K^+^ current [τ_decay_I_K_(5)/τ_decay_I_K_(1), **f**]. **g.** Glutamate transporter mediated current - astrocytic currents in response to stimulation of Schaffer collaterals after subtraction of K^+^ current: I_GluT_(1) – response to a single stimulus; I_GluT_(5) – isolated response to 5^th^ stimulus in burst stimulation (5 stim x 50 Hz minus 5 stim x 50 Hz). *Top* – adult, *bottom* – old mouse. **h,i.** Summary of activity-dependent change in the amplitude [I_GluT_(1)/I_GluT_(5), **h**] and the decay time of glutamate transporter current [τ_decay_I_GluT_(5)/τ_decay_I_GluT_(1), **i**]. *Grey circles –* adult *and green circles –* old mice. The data are presented as the mean ± SEM. N.S. *p* > 0.05; \**p* < 0.05; \*\**p* < 0.01; two-tailed two-sample *t*-test.

### 2.4. Ageing compromises K^+^ clearance and glutamate uptake

Decreased VF of fine astrocyte processes may associate with reduced astrocytic coverage of synapses, which, in consequence, may affect glutamate and K^+^ clearance (Oliet et al., 2001; Plata et al., 2018; Popov et al., 2020). To test this hypothesis, we monitored glutamate transporter and K^+^ uptake mediated currents (I_GluT_ and I_K_, respectively) in voltage-clamped CA1 *str.radiatum* astrocytes in response to stimulation of Schaffer collaterals. Because the kinetics of I_K_ is much slower than that of I_GluT_, we recorded the amplitude of I_K_ at 200 ms after the stimulus. We applied a single stimulus and two burst stimulations (4 stim x 50 Hz and 5 stim x 50 Hz). Then we subtracted the response to 4 stimuli from the response to 5 stimuli. The resulting response [I_K_(5)] was compared to the response to the single stimulus [I_K_(1)]. This ratio characterised activity-dependent changes in the I_K_ (Fig. 3d). No significant difference between two age groups was observed in I_K_(5)/I_K_(1) amplitude ratio (1.05 ± 0.15, *n* = 6, in the adult; 0.77 ± 0.11, *n* = 5, in the old; *p* = 0.09, two-sample *t*-test; Fig. 3e). However, the ratio of decay times τ_decay_I_K_(5)/τ_decay_I_K_(1) was significantly larger in the old animals (0.81 ± 0.13, *n* = 6, in the adult; 1.15 ± 0.10, *n* = 5, in the old; *p* = 0.049, two-sample *t*-test; Fig. 3f). This result points to the reduced efficiency of K^+^ clearance with ageing.

Next, we employed a similar experimental design to measure activity-dependent changes in the amplitude and decay time of I_GluT_. The I_GluT_ was recorded with blocked ionotropic glutamate receptors (see Method section), which are responsible for the most of K^+^ released during synaptic transmission (Lebedeva et al., 2018; Shih et al., 2013; Sibille et al., 2014). The residual I_K_, presumably mediated by K^+^ release associated with action potentials, was dissected at the end of each experiment with glutamate transporter blocker (100 μM TBOA). This I_K_ was scaled to the tail of currents recorded during the experiment and then subtracted from them resulting in pure I_GluT_ (Fig. 3g). No significant difference between two age groups was observed in I_GluT_(5)/I_GluT_(1) amplitude ratio (1.15 ± 0.09, *n* = 6, in the adult; 1.13 ± 0.07, *n* = 5, in the old; *p* = 0.86, two-sample *t*-test; Fig. 3h). Similarly to I_K_ the ratio of decay times τ_decay_I_GluT_(5)/τ_decay_I_GluT_(1) was significantly larger in the old animals (0.81 ± 0.09, *n* = 6, in the adult; 1.11 ± 0.12, *n* = 5, in the old; *p* = 0.04, two-sample *t*-test; Fig. 3i). This result indicates an age-dependent reduction in astrocytic glutamate uptake.

### 2.5 Ageing reduces spread of Ca^2+^ events in hippocampal astrocytes

Astrocytic Ca^2+^ activity plays important role in physiology of these cells (Semyanov, 2019; Verkhratsky and Nedergaard, 2018). It triggers secretion (Zorec et al., 2012), regulates local blood flow (Filosa et al., 2004; Marina et al., 2020), defines astrocyte morphological plasticity (Molotkov et al., 2013; Tanaka et al., 2013). On the other hand, astrocytic Ca^2+^ activity depends on astrocyte morphology and intercellular gap-junction coupling (Semyanov, 2019; Wu et al., 2019). To test whether astrocyte atrophy in the aged brain affects Ca^2+^ dynamics in the astrocytic network, we performed confocal Ca^2+^ imaging in CA1 *str.radiatum* of mouse hippocampal slices loaded with membrane-permeable Ca^2+^ dye Oregon-Green 488 BAPTA-1 AM (OGB-1). This dye predominantly stains astrocytes in slices however it can, potentially, report Ca^2+^ activity in neurones. Luckily, astrocytic Ca^2+^ events have different properties: they are longer then neuronal and have distinct spatial spread (Semyanov, 2019). Therefore, we analysed only Ca^2+^ transient that lasted longer than 4 s, assuming that they mostly have an astrocytic origin. Ca^2+^ transients were first detected in individual pixels, these active pixels were combined to obtain spatiotemporal Ca^2+^ events in three-dimensional space (x-y-time; Fig. 4a,b) (Wu et al., 2014). No significant difference in the frequency density of Ca^2+^ events was observed for two age groups (0.9 ± 0.3 s^−1^μm^−2^, *n* = 11 in the adult; 1.2 ± 0.7 s^−1^μm^−2^, *n* = 6 in the old; *p* = 0.32 two-sample *t*-test). Next, we analysed Ca^2+^ event duration, area (maximal projection along time axis into the x-y plane) and volume. The duration of Ca^2+^ events significantly increased in old animals (median 6 s, *n* = 358 [11 slices]; in the adult; 7 s, *n* = 248 [6 slices] in the old; *p* = 0.005, Mann-Whitney test; Fig.4c), while the area decreased (median 20.2 μm^2^, *n* = 358 [11 slices] in the adult; 16.2 μm^2^, *n* = 248 [6 slices] in the old; *p* = 0.02, Mann-Whitney test; Fig.4d). These two changes compensated each other in terms of overall Ca^2+^ thus making the event volume similar (median 57.8 s*μm^2^, *n* = 358 [11 slices] in the adult; 51.4 s*μm^2^, *n* = 248 [6 slices] in the old; *p* = 0.33, Mann-Whitney test; Fig.4e). We also noticed that prolongation of Ca^2+^ events in the old mice could be explained by merging of subsequent smaller events because of multiple events initiation at the same spot. Therefore, we analysed Ca^2+^ event initiation spots and discovered that events less frequently generated at the same spot in the adult than in the old mice (Fig. 4f,g). Hence, astrocytic Ca^2+^ events in the adult brain have more stochastic nature in the adult brain and become linked to spatially restricted Ca^2+^ microdomains in the old.

**Figure 4.**
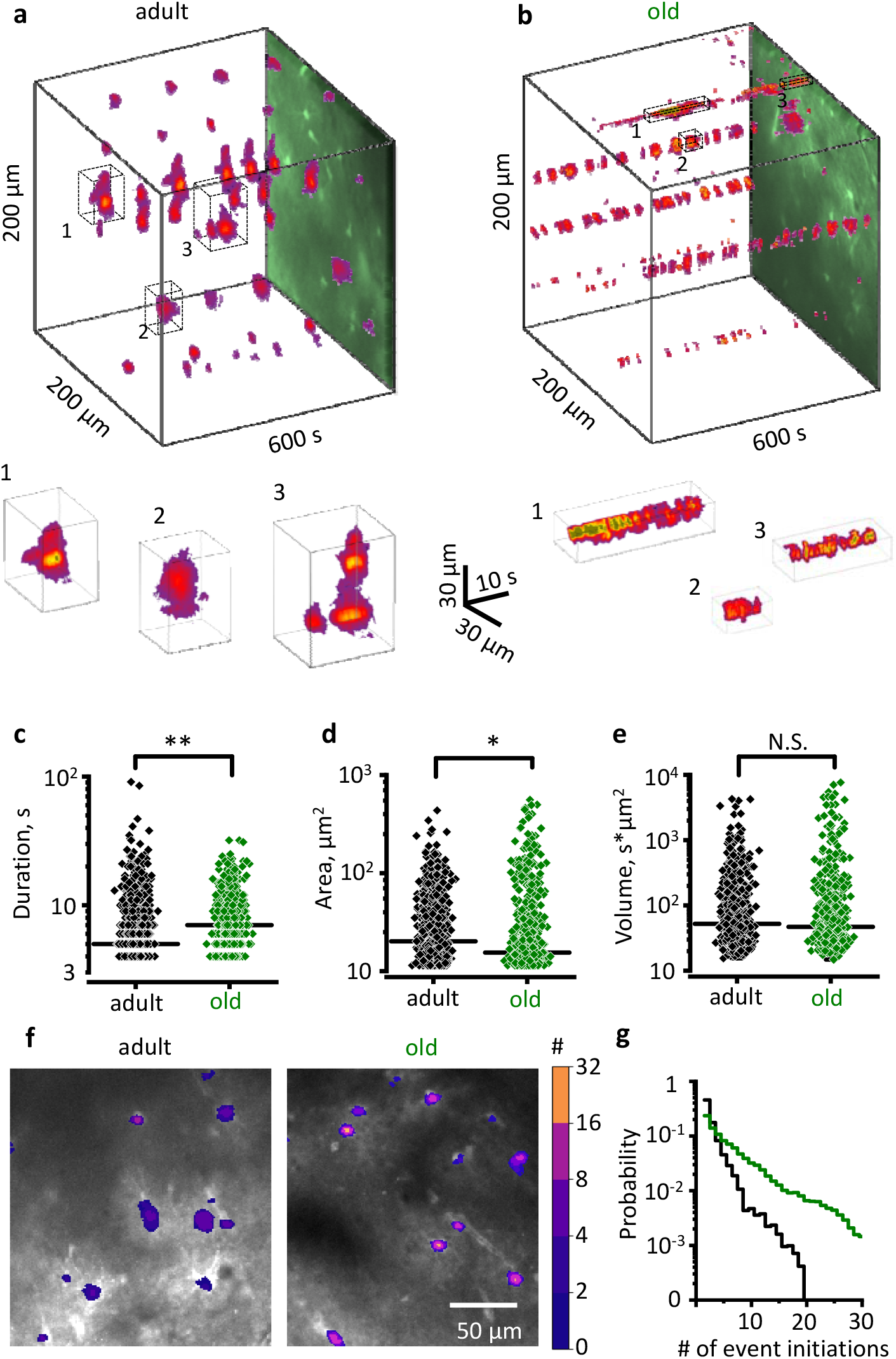
Ageing reduces spread of Ca^2+^ events in hippocampal astrocytes. **a,b.** x-y-time reconstruction of detected Ca^2+^ events in hippocampal *str.radiatum* (*top*) with examples of individual events zoomed in from the boxed regions (*bottom*) in the adult (**a**) and old (**b**) mice. **c-e.** Summary of event duration (**c**), area (maximal projection, **d**), and volume (**e**) in the adult (black diamonds) and old (green diamonds) mice. Horizontal black bar indicates median. **f.** Ca^2+^ event initiation points with colour-coded number of events initiated during 10 min recording. **g.** The distribution of Ca^2+^ events initiation points according the number of events initiated per pixel in the adult (black trace) and old (green trace) mice. \**p* < 0.05; \*\**p* < 0.01; Mann-Whitney test.

### 2.6. Reduced astrocytic coverage of synapses parallels impaired synaptic plasticity

Decreased glutamate uptake leads to glutamate spillover which activates extrasynaptic NR2B-containing NMDA receptors (Nie and Weng, 2010; Scimemi et al., 2004). This, in turn, can reduce the magnitude of hippocampal long-term potentiation (LTP) (Valtcheva and Venance, 2019). Therefore, we tested if astrocyte dystrophy in the old age is also associated with impaired LTP. We recorded field excitatory postsynaptic potential (fEPSP) with extracellular glass electrode in CA1 *str.radiatum* of mouse hippocampal slices in response to extracellular stimulation of Schaffer collaterals with a bipolar electrode located in *str.radiatum* at the border of CA1 and CA2. LTP was induced with high-frequency stimulation (HFS, 20 stim. x 100 Hz repeated 3 times with 20-s intervals) and estimated as a change of fEPSP amplitude in percentage relatively to pre-HFS baseline fEPSP amplitude. In agreement with previous publications reduced glutamate clearance correlated with impaired LTP in the old animals (LTP magnitude was 153 ± 4 % of baseline, *n* = 7, in the adult; and 128 ± 4 % of baseline, *n* = 6, in the old; *p* = 0.001, two-sample *t*-test; Fig. 5).

**Figure 5.**
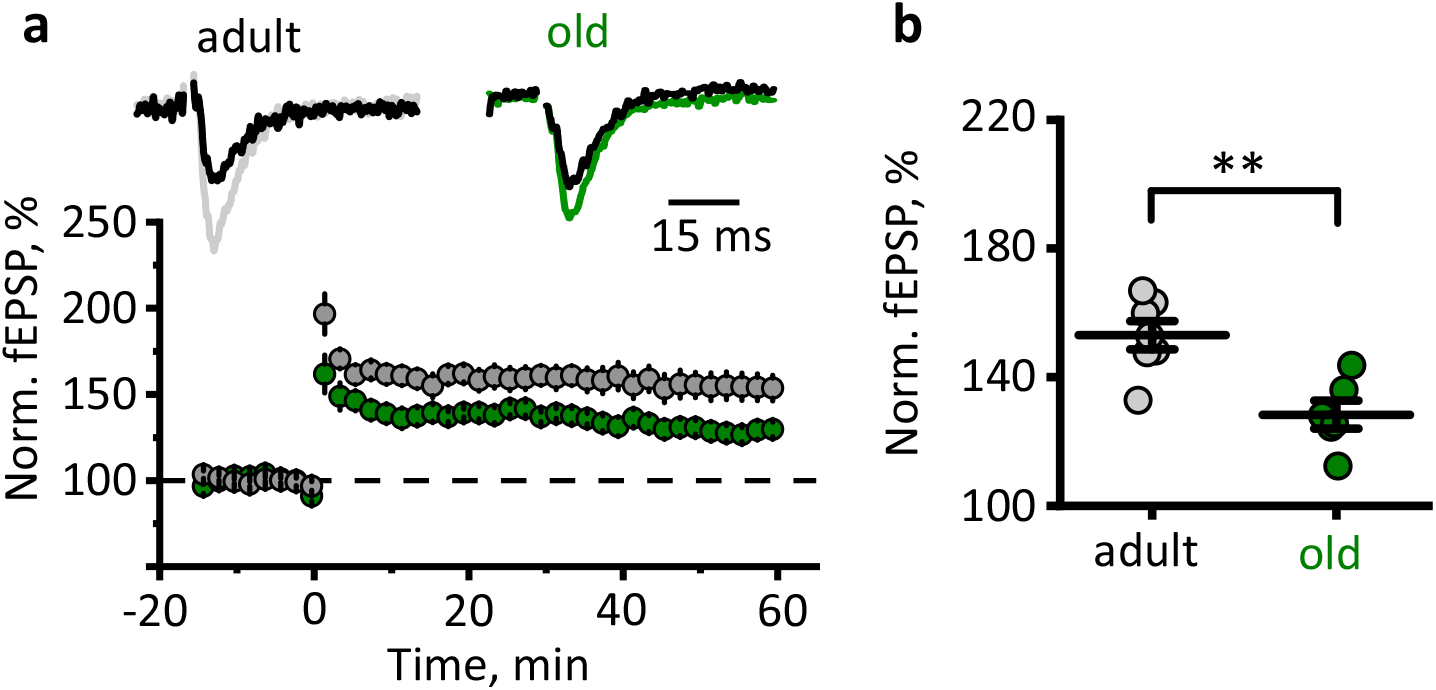
Reduced LTP in CA3-CA1 synapses of old mice. **a.** Timecourse of fEPSP amplitude recorded in CA1 *str.radiatum* in response to Schaffer collaterals stimulation before and after HFS in the adult and old mice hippocampal slices. Insets show sample fEPSP before and 60 min after HFS. 0 min point is the time of HFS. fEPSP amplitude is normalised to mean fEPSP amplitude prior to HFS. **b.** Summary of LTP magnitude 50-60 min after HFS stimulation. *Grey circles –* adult *and green circles –* old mice. The data are presented as the mean ± SEM. \**p* < 0.05 two-tailed two-sample *t*-test.

## 3. Discussion

### 3.1. Shrinkage of astrocytic territorial domains in the old age

The concept of increased astrocytic reactivity and astrocytic hypertrophy in the aged brain is widespread (Lynch et al., 2010; Salas et al., 2020; Unger, 1998). This concept associates with views on progressive activation of microglia in ageing (Norden and Godbout, 2013). The notion of generalised gliosis in the senescent brain is linked to a more global theory of “inflammageing” (Franceschi, 2007) that regards the ageing process as a mounting chronic inflammation. These conceptualisations, however, are based on relatively thin data when it comes to astrogliosis; the ubiquitous gliosis is further questioned by well documented presence of dystrophic microglia in the aged human brains (Conde and Streit, 2006). There are substantial arguments for a decrease in functional capacities of neuroglial cells in ageing, with glial dystrophy and paralysis facilitating neurodegeneration (Streit et al., 2020; Verkhratsky et al., 2015; Verkhratsky et al., 2019).

Changes in GFAP-positive astrocytic profiles are not universally observed throughout the brain in aged animals (Rodriguez-Arellano et al., 2016); and moreover, even an increase in GFAP-labelled astrocytic cytoskeleton does not necessarily translate into cellular hypertrophy: the volume of reactive astrocytes undergoing anisomorphic astrogliosis does not change (Wilhelmsson et al., 2006). Furthermore, an increase in GFAP-stained cytoskeleton may even develop in parallel with atrophy of peripheral astrocytic processes as has been demonstrated in experimental epilepsy (Plata et al., 2018). The data accumulated on astrocytic ageing so far do not unequivocally support the notion of increased astrocytic reactivity in the senescent brain.

Analysis of GFAP morphological profiles misses peripheral processes altogether because they do not express this intermediate filament. At the same time, peripheral processes account for up to 80% of the total surface area of a typical protoplasmic astrocyte (Verkhratsky and Nedergaard, 2018). These membranous leaflets represent functionally important compartment, which provides for perisynaptic coverage and forms astrocytic cradle (Gavrilov et al., 2018; Verkhratsky and Nedergaard, 2014); peripheral processes also establish contacts with other astrocytes and make astrocytic syncytia. Here we addressed the question of age-dependent changes in fine astrocytic morphology by analysing high resolution two-photon optical sections of astrocytes filled with low molecular weight probe Alexa Fluor 594. This fluorescent dye diffuses through the cytosol and penetrates in the most distant parts of the cell. In particular, the signal of Alexa 594 from tiny, optically unresolved, processes contribute to a VF of astrocytic profile thus allowing quantification of these processes. Two-photon imaging, in conjunction with 3D reconstruction, Sholl analysis and VF measurements, demonstrated a significant reduction in astrocytic profiles and in astrocytic territorial domains in aged brains. Ageing did not affect the total number of astrocytes; neither the numbers of primary astrocytic branches has been changed. Ageing, however, significantly reduced complexity of arborisation (as judged by the reduction in the number of intersections, Fig. 1d), length of processes (Fig. 1e) and the domain area (Fig. 1g). The shrinkage of the domain area was quite substantial: it reduced from 800 μm^2^ to ~ 500 μm^2^. We also found a considerable decrease of VF associated with peripheral astrocytic processes or leaflets – it was almost halved in the old astrocytes (Fig. 2b). Collectively, these changes reduce astrocytic contribution to the neuropil and can potentially widen diffusion channels in the brain interstitium. Our results, therefore, provide a cellular basis for an increase in mean diffusivity of the grey matter in of humans observed with diffusion tensor imaging (Salminen et al., 2016). In addition, astrocyte shrinkage without changes in the cell density inevitably sets the astrocytic domain borders apart from each other. This, in turn, reduces astrocyte-to-astrocyte connections, which we have observed as reduced intercellular dye diffusion through gap-junctions, corroborating attenuated astrocyte coupling.

### 3.2. Age-dependent remodelling of astrocytic Ca^2+^ signalling

Changes in Ca^2+^ homeostasis and Ca^2+^ signalling has been considered as a general mechanism of ageing of neural cells, which was formalised in a “calcium hypothesis of ageing” (Khachaturian, 1987; Verkhratsky and Toescu, 1998). In developing this hypothesis, the neurodegenerative diseases (that are often regarded as somewhat exacerbated ageing) have been considered as “calciumopathies” (Stutzmann, 2007). More detailed analysis, however, revealed that physiological ageing is associated with rather subtle modifications of Ca^2+^ homeostatic machinery in neurons, which renders them more susceptible to long-lasting excitotoxic challenges; in contrast, pathological ageing with neurodegeneration is indeed associated with aberrant Ca^2+^ signalling (Busche and Konnerth, 2016; Elena et al., 2020; Toescu and Verkhratsky, 2007; Toescu et al., 2004). We found a somewhat peculiar spatial reorganisation of astrocytic Ca^2+^ signals in aged animals. The stochastic distribution of Ca^2+^ signals in the adult mice has been replaced with a relatively rigid link of Ca^2+^ microdomains to fixed locations. Arguably, shrinkage of astrocytes may be inadequate to support the variability of subcellular Ca^2+^ event initiation points. Localisation of Ca^2+^ events may also reflect reduced astrocyte-to-astrocyte connectivity that hampers the propagation of Ca^2+^ signals. Whichever the case, this finding is consistent with a general decrease in overall plasticity in the ageing brain, which relies on pre-existing signalling routes being reluctant or unable to establish new ones.

### 3.3. Reduced astrocytic synaptic coverage parallels impaired synaptic plasticity

The decrease in astrocytic territorial domains and general shrinkage of astrocytes may have numerous consequences for the nervous tissue. In particular, atrophy of perisynaptic processes can substantially modify synaptic transmission. Astrocytes, through a wide array of homeostatic transporters localised in the perisynaptic membrane (Verkhratsky and Rose, 2020), control ionostasis and neurotransmitters in the synaptic cleft and supply neurons with neurotransmitter precursors and energy substrates (Verkhratsky and Nedergaard, 2014). Retraction of astrocytic processes and reduction in astrocytic synaptic coverage can, therefore, impact on synaptic transmission and synaptic plasticity. We found that atrophic old astrocytes cannot support K^+^ buffering and uptake of glutamate as effectively and astrocytes in the adult brain. This failure in homeostatic support translates into deficient LTP, which most likely results from an increased spillover of glutamate and activation of extrasynaptic NMDA receptors (Valtcheva and Venance, 2019). Our results, therefore, indicate that age-dependent impairment of learning and memory originates from impaired synaptic plasticity associated with astrocyte shrinkage and retraction of perisynaptic astrocytic processes giving way to glutamate spillover.

### 3.4. Recapitulation

In summary, we provide compelling evidence that physiological ageing is associated with morphological atrophy of astrocytic peripheral branchlets and leaflets. This morphological decline is associated with (and most likely is a cause of) functional asthenia of astrocytes manifested by diminished efficacy of glutamate and K^+^ uptake, and remodelling of Ca^2+^ activity towards more localised patterns. This functional decline of astrocytes favours glutamate spillover which is arguably responsible for decreased synaptic plasticity in the senescent brain.

## 4. Material and methods

### 4.1. Animals

All procedures were done in accordance with the Shemyakin-Ovchinnikov Institute of bioorganic chemistry ethical regulations. The experiments were performed in C57BL/6 male mice of three age groups: 3-4 months old (young), 9-12 months old (adults) and 20-24 months (old).

### 4.2. Slice preparation

The mice were anaesthetized with isoflurane (1-chloro-2,2,2-trifluoroethyl-difluoromethyl ether) before being sacrificed. Intracardiac perfusion 5 ml/min was done before slice preparation. The perfusion solution contained (in mM): 92 *N*-methyl-D-glucamine (NMDG), 2.5 KCl, 1.25 NaH_2_PO_4_, 30 NaHCO_3_, 20 HEPES, 25 glucose, 2 thiourea, 5 Na-ascorbate, 3 Na-pyruvate, 0.5 CaCl_2_ꞏ2H_2_O, and 10 MgSO_4_ꞏ7H_2_O. After perfusion, the brain was exposed and placed in in the same solution which was used for perfusion. Hippocampi were dissected and cut into transverse slices (350 μm) with a vibrating microtome (Microm HM650 V; Thermo Fisher Scientific). Slices were left for extended recovery and ionic equilibration for 2.5 hours (30 min at 32-34°C and then for 2 hours at room temperature) in a storage solution containing (in mM): 127 NaCl; 2.5 KCl; 1.25 NaH_2_PO_4_; 2 MgCl_2_; 1 CaCl_2_; 25 NaHCO_3_; 25 D-glucose. The experiments were carried out at 34°C in immersion chamber with continuous superfusion (1-3 ml/min) by solution containing (in mM): 127 NaCl; 2.5 KCl; 1.25 NaH_2_PO_4_; 1 MgCl_2_; 2 CaCl_2_; 25 NaHCO_3_; 25 D-glucose. All solutions had an osmolarity of 295 ± 5 mOsm and a pH of 7.4 and were saturated with carbogen (95% O_2_ and 5% CO_2_).

### 4.3. Astrocyte morphometry

Astrocytes were loaded with fluorescent dye Alexa Fluor 594 through the patch pipette which was used for electrophysiological recordings (see below). Images were collected with W Plan-APOCHROMAT 40x/1.0 water immersion objective, using the Zeiss LSM 7 MP microscope (Carl Zeiss, Germany) coupled with Ti:sapphire femtosecond laser Chameleon Vision II (Coherent, UK).

### 4.4. 3D Sholl analysis

All processing steps were performed using image-funcut library [image-funcut, https://github.com/abrazhe/image-funcut] and other custom-written Python scripts, using Scikit-Image [scikit, http://scikit-image.org/] and Sci-Py [scipy, http://www.scipy.org/] libraries.

In brief, Z-stacks corresponding to the emission spectrum (565-610 nm) of Alexa Fluor 594 (resolution was 512×512 pixels (0.2 μm/px) on XY axis with a step on Z-axis 1 μm/frame 100 frames in total) were re-sampled to the same lateral resolution of 0.2 μm/px. Coherence-enhancing diffusion filtering was performed for each image in the stack to enhance all filamentous structures.

Before further analysis, estimates of PMT gain and offset values were made to convert PMT values to photon count estimates. To this end, small cubic patches from the stack were randomly selected, and the dependence of sample variance vs sample mean fitted by a straight line, with the slope corresponding to PMT gain and X-axis intercept corresponding to PMT offset.

This renormalization was needed for correct astrocyte volume fraction analysis.

Binary masks representing astrocyte processes were created for 3D Sholl analysis. Prior to binarization, elongated structures corresponding to astrocytic processes were emphasized by applying coherence enhancing diffusion filtering to all planes along each axis and averaging the result (Weickert and Scharr, 2002). Binarization was done by hysteresis thresholding with a low and high threshold corresponding to 1x and 3x standard deviations of noise in the analysed stack, followed by pruning all structures less than 100 voxels.

After binarization, we created a 3D map of branches and branchlets for each cell by using a simple neurite tracer plugin in ImageJ. We constructed concentric spheres with increasing radius (from 5 to 50 μm with1 μm step) from the centre of the astrocyte soma and extracted different parameters by intersection spheres with branch map for each cell. The domain area of each astrocyte was determined as the area occupied by astrocyte processes (maximum projection).

### 4.5. Estimation of volume fraction (VF)

An image containing astrocyte soma was selected in the z-stack. Special attention was paid that the fluorescence of soma was not saturated (Popov et al., 2020; Wu et al., 2019). Five cross-sections for each cell were plotted via the centre of soma at the angle of 72° from each other. We get points from two sides from each cross-section and obtained 10 FV values for each cell.

The fluorescent profiles along these lines were obtained, and large fluctuations (> 10% and >0.5 μm) of fluorescence corresponding to astrocytic branches and branchlets were cut out. VF was estimated as fluorescence intensity in unresolved processes normalized fluorescence intensity in soma. Characteristic VF was obtained at distance 10-20 μm from the soma border (supplementary Fig. 1).

### 4.6. Electrophysiological recordings

Neuronal and astrocytic responses were evoked by extracellular stimulation of the Schaffer collaterals with a bipolar stimulating electrode (FHC, Bowdoinham, USA) placed in the *str.radiatum* at the CA1–CA2 border. The stimulation was performed with rectangular current pulses (duration: 0.1 ms, interval: 20 s) with DS3 isolated current stimulator (Digitimer Ltd, UK). Responses were amplified with a Multiclamp 700B amplifier (Molecular Devices, USA), digitized with digital-analogue converter board NI PCI-6221 (National Instruments, USA), and recorded with WinWCP v5.2.3 software by John Dempster (University of Strathclyde). The data were analysed with the Clampfit 10.2 software (Molecular Devices, USA).

### 4.7. Field potential recordings and LTP induction

The field excitatory postsynaptic potentials (fEPSPs) were recorded in CA1 *str.radiatum* with glass microelectrodes (resistance: 2–5 MOhm). For timecourse experiments, half-maximal stimulus intensity was chosen (the stimulus intensity when fEPSP amplitude was in 40–50% of the amplitude when the population spike appeared). The strength of stimulation was constant during the experiment, usually being 100–150 μA. The LTP was induced if the stable amplitude of the baseline fEPSP could be recorded for 15 min. Three trains of high-frequency stimulation (HFS, 20 pulses at 100 Hz, with an inter-train interval of 20 s protocol) were applied to induce LTP. The fEPSPs were recorded after induction protocol for at least 60 min. The LTP magnitude was estimated as the ratio of potentiated fEPSP amplitude (averaged in the interval of 50-60 min after the HFS) to baseline fEPSP amplitude.

### 4.8. Astrocytic recordings

Astrocytes were selected in the *str.radiatum* at a distance of 100 – 200 μm from the stimulating electrode and whole-cell recorded with borosilicate pipettes (3 - 5 MΩ) filled with an internal solution containing (in mM): 135 KCH_3_SO_3_, 10 HEPES, 10 Na_2_phosphocreatine, 8 NaCl, 4 Na_2_-ATP, 0.4 Na-GTP (pH adjusted to 7.2 with KOH; osmolarity to 290 mOsm). 50 μM Alexa Fluor 594 (Invitrogen, USA) was added to the internal solution for morphological study. Passive astrocytes were identified by their small soma (5 – 10 μm diameter), strongly negative resting membrane potential, and linear current-voltage (I-V) relationship. In current-clamp mode, current steps were applied to corroborate the absence of membrane excitability. In voltage-clamp recordings the astrocytes were held at − 80 mV. Voltage steps (Δ20mV; 500 ms) from - 140mV to + 80 mV were applied to obtain I-V relationships.

Input resistance was calculated according to the Ohm law by dividing voltage step magnitude (−5 mV) by current response.

One, four and five electrical stimuli at 50 Hz were applied to Schaffer collaterals to induce synaptic currents in the astrocytes followed by a voltage step of −5 mV for monitoring cell input resistance. Signals were sampled at 5 kHz and filtered at 2 kHz. Astrocyte currents in response to one, four, and five stimuli were baseline subtracted and averaged respectively. Current in response to the fifth stimulus was obtained by the subtraction of current evoked by four stimuli from current evoked by five stimuli.

### 4.9. Analysis of potassium current (I_K_) and glutamate transporter current (I_GlutT_)

The astrocytic currents induced by synaptic stimulation represent a sum of several components including local filed potential mediated current, fast glutamate transporter current (I_GluT_) and slow K^+^ inward current (I_K_) (Sibille et al., 2014). I_K_ lasts for hundreds of ms and, hence, was analysed 200 ms after the last stimulus, where it is not contaminated by the other current. The amplitude of the I_K_ was measured at this time point. The I_K_ decay was fitted with a mono-exponential function, and τ_decay_ was calculated.

I_GluT_ was obtained in the presence of antagonists of postsynaptic ionotropic receptors: 25 μM NBQX – AMPA receptor antagonist, 50 μM D-APV – NMDA receptor antagonist, and 100 μM picrotoxin – GABA_A_ receptor channel blocker (all from Tocris Bioscience, UK). This prevented filed potential mediated component of astrocytic current and largely reduced I_K_ (Shih et al., 2013). After 10 minutes of recordings, 100 μM DL-TBOA (Tocris Bioscience, UK), an excitatory amino acid transporter (EAATs) blocker, was added to the bath to obtain residual I_K_ which was fit to the tail and subtracted from synaptic current to obtain pure I_GluT_. The I_GluT_ decay was fitted with mono-exponential function and τ_decay_ was calculated.

### 4.10 Ca^2+^ imaging

Ca^2+^ activity was recorded with a confocal microscope, Zeiss LSM DuoScan 510, in CA1 *str.radiatum* of hippocampal slices pre-incubated with Ca^2+^ dye, OGB-1 (Invitrogen, USA) and an astrocyte-specific marker, sulforhodamine 101 (100 nM, Invitrogen, USA). After the preparation, the slices were transferred to a 3 ml incubation chamber with constantly gassed storage solution containing both dyes. OGB-1 was initially dissolved to 0.795 mM in 0.8% Pluronic F-127 in DMSO. Then 3 μl of the dye was added to the chamber. After incubation for 40 - 45 min at 37 °C in the dark, the slices were transferred to the recording/imaging chamber for time-lapse imaging (one frame/s). OGB-1 was excited with a 488 nm argon laser and imaged with an emission band-pass filter 500 - 530 nm; sulforhodamine 101 was excited with a 543 nm HeNe laser and imaged with an emission band-pass filter 650 - 710 nm. The imaging was performed for 10 min at 34 °C in normal ASCF, then 30 dark noise images were recorded.

## Aknowlegements

The research was supported by Russian Science Foundation grant 20-14-00241

## Supplementary information

**Supplementary Figure 1.**
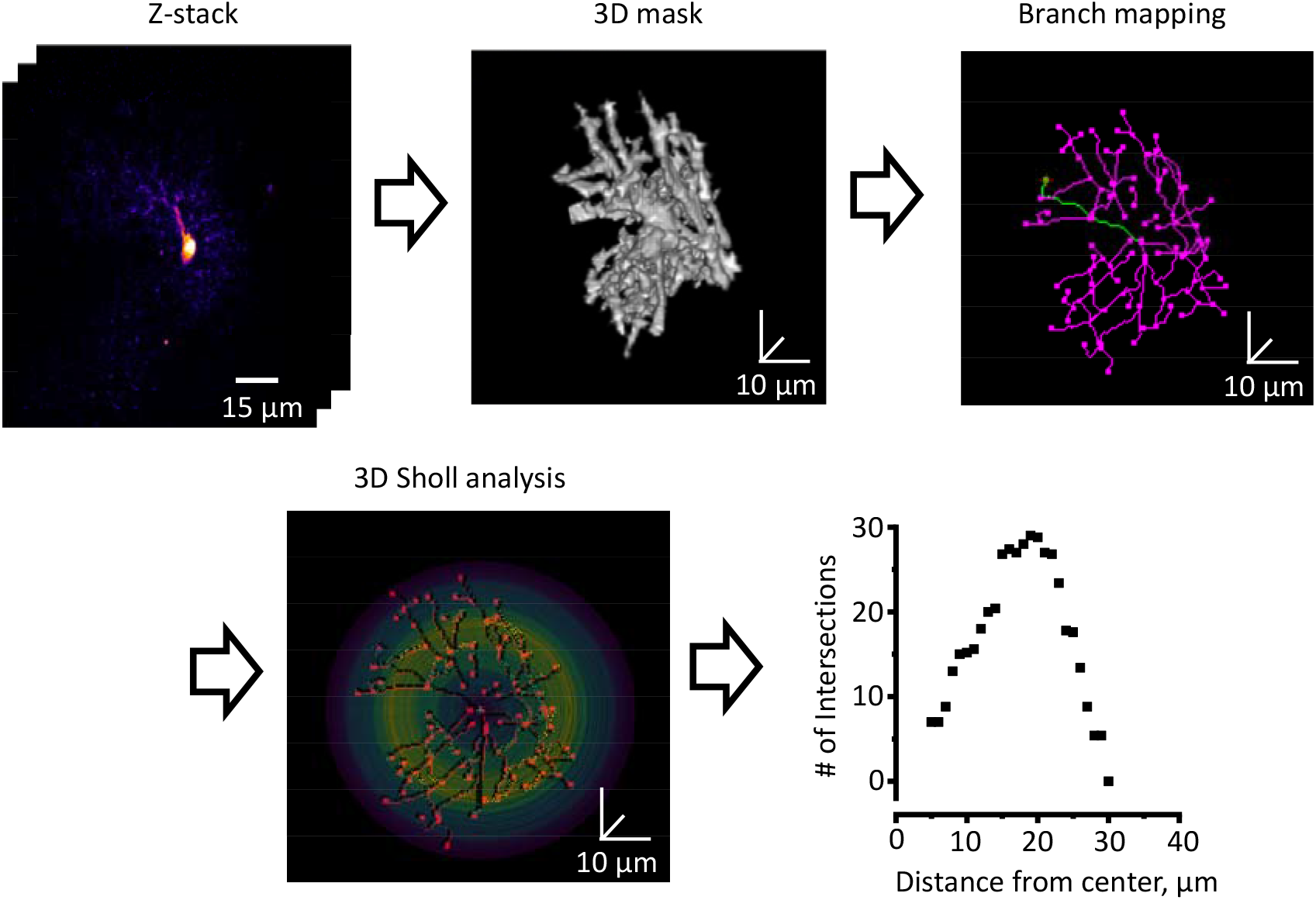
Method of 3D Sholl analysis. First, Z-stack of confocal images of astrocytes loaded with fluorescent dye, Alexa Flo 594, through patch pipette was obtained. Second, three-dimensional (3D) mask of reconstructed astrocyte was obtained with custom-written Python script. Third, the branch and branchlet mapping along the mask was performed in ImageJ (https://imagej.net/). The 3D Sholl analysis was performed with concentric spheres centered in the middle of astrocyte soma. The number of intersections of the branches and branchlets with the spheres was estimated.

