## Supplementary information for "Astrocytes dystrophy in ageing brain parallels impaired synaptic plasticity"

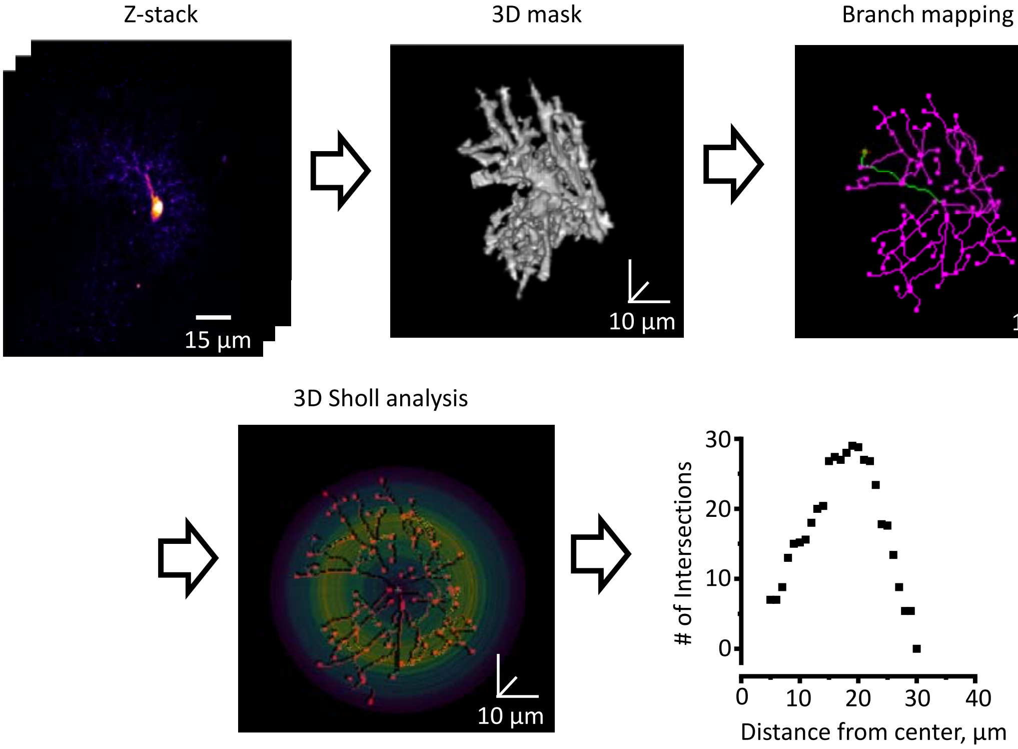


**Supplementary Figure 1 Method of 3D Sholl analysis**

First, Z-stack of confocal images of astrocytes loaded with fluorescent dye, Alexa Flo 594, through patch pipette was obtained. Second, three-dimensional (3D) mask of reconstructed astrocyte was obtained with custom-written Python script. Third, the branch and branchlet mapping along the mask was performed in ImageJ (https://imagej.net/). The 3D Sholl analysis was performed with concentric spheres centered in the middle of astrocyte soma. The number of intersections of the branches and branchlets with the spheres was estimated.
